# Brain aromatase and its relationship with parental experience and behavior in male mice

**DOI:** 10.1101/2023.07.15.549144

**Authors:** Paula Duarte-Guterman, Dimitri A. Skandalis, Ariane Merkl, Diana B. Geissler, Günter Ehret

## Abstract

In most mammals, paternal care is not mandatory for raising offspring. In house mice, experience with pups governs the extent and quality of paternal care. First-time fathers undergo a dramatic transition from ignoring or killing pups to caring for pups. The behavioral shift occurs together with changes in brain estrogen signaling as indicated by changes in estrogen receptor presence and distribution in multiple areas regulating olfaction, emotion, and motivation. Here, we report changes in the expression of aromatase, the enzyme converting testosterone into estrogen, as an indirect measure of estrogen synthesis. The amount of paternal experience (5 or 27 days) was associated with increased numbers of immunocytochemically-identified aromatase expressing cells in the medial and cortical amygdala, posterior piriform cortex, and ventromedial hypothalamus. Functionally, these changes can be related to the disappearance of aggression or neglect towards pups when first-time fathers or, even more, well-experienced fathers are handling their own pups. In the lateral septum, the anterior piriform cortex and to some extent in the medial preoptic area, parental experience increased the number of aromatase-positive cells only in fathers with 27 days of experience, and only in the right hemisphere. This is a new case of brain-functional lateralization due to experience that has activated certain instinctive behavior. Nuclei/areas associated with maternal care (medial preoptic area, bed nucleus of stria terminalis, nucleus accumbens) exhibited a left-hemisphere advantage in aromatase expressing cells, both in pup-naïve and pup-experienced males. This newly found lateralization may contribute to the left-hemisphere dominant processing and perception of pup calls to release parental behavior. In general, the experience-dependent changes in aromatase expression we observed in most brain areas did not mirror the previously reported changes in estrogen receptors (ERα) when pup-naïve males became pup-caring fathers. Hence, paternal behavior may depend in a brain area-specific way on the differential action of estrogen through its receptors and/or direct local modulation of neural processing.

## Introduction

Mammalian paternal care is facultative, and the behavioral and neural dynamics that result in becoming a caring father are less well understood compared to those of obligatory maternal care (e.g., Bales and Saltzman, 2016; Brunton and Russell, 2008; Dulac et al., 2014; Guoynes and Marler, 2020; Horrell et al., 2021; Lonstein and De Vries, 2000; Numan, 2020; Numan and Insel, 2003; Rilling and Mascaro, 2017; Woodroffe and Vincent, 1994)). In rodents and primates, it is hypothesized that maternal care is induced by hormonal changes during pregnancy and parturition (onset phase) whereas the continuation of pup-caring (maintenance phase) is governed by sensory stimuli from the pups (Brunton and Russell, 2008; Fahrbach and Pfaff, 1982; Rosenblatt and Siegel, 1981). In house mice, pup-naïve females (i.e., having no sexual or pup caring experience), males, and even previous fathers do not experience the onset phase. Rather, they require various amounts of experience with pups (from hours up to several days of contact) for the activation of caring behavior (Akther et al., 2021; Alsina-Llanes et al., 2015; Ehret and Buckenmaier, 1994; Ehret and Koch, 1989; Ehret et al., 1987,1993; Elwood, 1985; Koch and Ehret, 1989; Martín-Sánchez et al., 2015; Stolzenberg and Rissman, 2011; Tachikawa et al., 2013). Nonetheless, pup-naïve females engage in consummatory (pup retrieval) and appetitive (preference for pup ultrasounds) behaviors (Koch and Ehret, 1991; Numan, 2006) after fewer days of experience with pups (and sometimes spontaneously; Alsina-Llanes et al., 2015; Martín-Sánchez et al., 2015) compared to either pup-naïve males or previous fathers (Ehret and Buckenmaier, 1994; Ehret et al., 1987,1993). Pup naïve females thus appear to be more easily sensitized or motivated by pup cues than males and even fathers to provide parental care. This observation is suggestive of sex differences in the neural mechanisms underlying the activation of maternal versus paternal instincts.

Numerous studies have characterized the role of the limbic regions in the activation and regulation of parental behaviors. These regions include the medial preoptic area (MPOA), bed nucleus of the stria terminalis (BNST), lateral septum (LS), nucleus accumbens (NAC), nuclei of the amygdala, and ventromedial hypothalamus (VMH). Olfaction is thought to provide the main modulatory sensory input, through pathways including the vomeronasal organ to the BNST and amygdala, the olfactory bulb to the amygdala, via the VMH to the LS, and via the piriform cortex (PIR) and hippocampus to the LS (e.g. Bridges, 2015; Brunton and Russell, 2008; Dulac et al., 2014; Ehret et al., 1991; Kohl and Dulac, 2018; Kohl et al., 2017; Numan, 2006, 2020). Estradiol treatment can activate both maternal and paternal instincts (Ehret and Koch 1989; Koch and Ehret, 1989; Poindron et al., 1988; Romero-Morales et al., 2018; Rosenblatt and Ceus, 1998; Rosenblatt et al., 1994; Siegel and Rosenblatt, 1975) and changes in brain estrogen signaling (through estrogen receptors) may be part of the underlying mechanism. In house mice, expression of parental behaviors coincides with changes in the distribution of estrogen receptor (ERα) positive cells in limbic brain areas of pup-naïve and pup-experienced females (Ehret and Buckenmaier, 1994; Koch, 1990) and males (Ehret et al., 1993). In pup-naïve male and female voles, high levels of spontaneous pup care are correlated with higher number of ERα positive cells in the MPOA (Li et al., 2015), potentially indicating convergent circuit dynamics.

Low levels of circulating estrogens in males (Nilsson et al., 2015) suggest that brain-active estrogens are primarily synthesized locally in the brain from testosterone by the enzyme aromatase (e.g., Cornil et al., 2006,2013; Garcia-Segura, 2008). Locally synthesized estrogens have been implicated in the onset of paternal behaviors in California mice (Trainor and Marler, 2002; Trainor et al., 2003). Aromatase is expressed in the limbic system of male mice (LS, BNST, MPOA, nuclei of the amygdala, VMH, and PIR) (Foidart et al., 1995; Schleicher et al., 1986; Stanić et al., 2014; Wozniak et al., 1992). In the present study, we sought to examine changes in aromatase expression during the onset and maintenance of paternal behavior of male mice (pup-naïve and fathers with increasing experience). We focused on LS, BNST, MPOA, nuclei of the amygdala, VMH, and PIR, which are areas in which ERα positive cells have also been located in pup-experienced fathers (Ehret et al., 1993).

## Materials and methods

### Animals

Adult male and female laboratory mice (outbred strain NMRI, 2-3 months old) with no previous sexual or parental experience were housed at the University of Ulm. As previously (Ehret et al., 1993), mice were randomly assigned to one of three treatment groups consisting of (1) naïve males without any experience with pups (N); (2) males co-caring with their female mate for their first litter for five days (5D); or (3) males co-caring with their female mate for their first litter until weaning and their second litter for five days. The total co-caring time of the latter males (first and second litter) was 26-27 days (27D) since after delivery of the first litter, females in post-partum estrus paired immediately and, while nursing the first litter up to weaning at 21 days, delivered the second litter after 21-22 days of gestation. Mice were housed at 22°C and a 12 h light-dark cycle (starting at 7.00 h) in plastic cages (26.5 × 20 × 14 cm) either with other naïve males (group N; 4 brothers per cage) or with one female (groups 5D and 27D). Water and food were available *ad libitum*. The experiments were carried out in accordance with the European Communities Council Directive (86/609/EEC) and were approved by the appropriate authority (Regierungspräsidium Tübingen, Germany).

### Pup retrieval tests

Testing of pup retrieval behavior was performed as previously described (Ehret and Koch, 1989; Ehret et al., 1993). Male mice (alone or with litter mate and pups) were placed overnight on a running board (length 110 cm, width 8 cm) with a central nest depression (Fig. 1A) suspended in a sound-proof and anechoic room. The running board was supplied with mouse chow and nest material from the cage and covered with a plexiglass hood having an opening for the tip of a suspended water bottle. The next day, retrieving tests were done under dim red light (<1 lx) ensuring that males relied on the sense of smell, audition, and touch to respond and retrieve pups. Before the start of testing, the hood, and the female with the pups (groups 5D and 27D) were removed and placed back to their home cage outside the soundproof room. Seven pups were kept in the room to serve as targets for retrieving. The males were habituated for at least 30 min to the new situation after the hood (group N) or the hood and female with pups (groups 5D and 27D) were removed. Prior to testing males of the N group, five-day old foster pups were rubbed with nest material of the respective males to ensure a familiar scent. During the test, the seven pups (always 1-3 pups at a time) were randomly placed on the running board, at least 30 cm away from the nest. A male was scored as ‘retriever’ if he retrieved all seven pups to the nest within 10 min. A male was scored as ‘non-retriever’ if he had not retrieved any pups after 10 mins. The distinction between retriever and non-retriever was clear-cut, because once a male had retrieved the first pup, he retrieved all. Retrieving tests were immediately stopped and the pups removed if a male attacked any of the pups (‘aggressor’).

**Figure 1.**
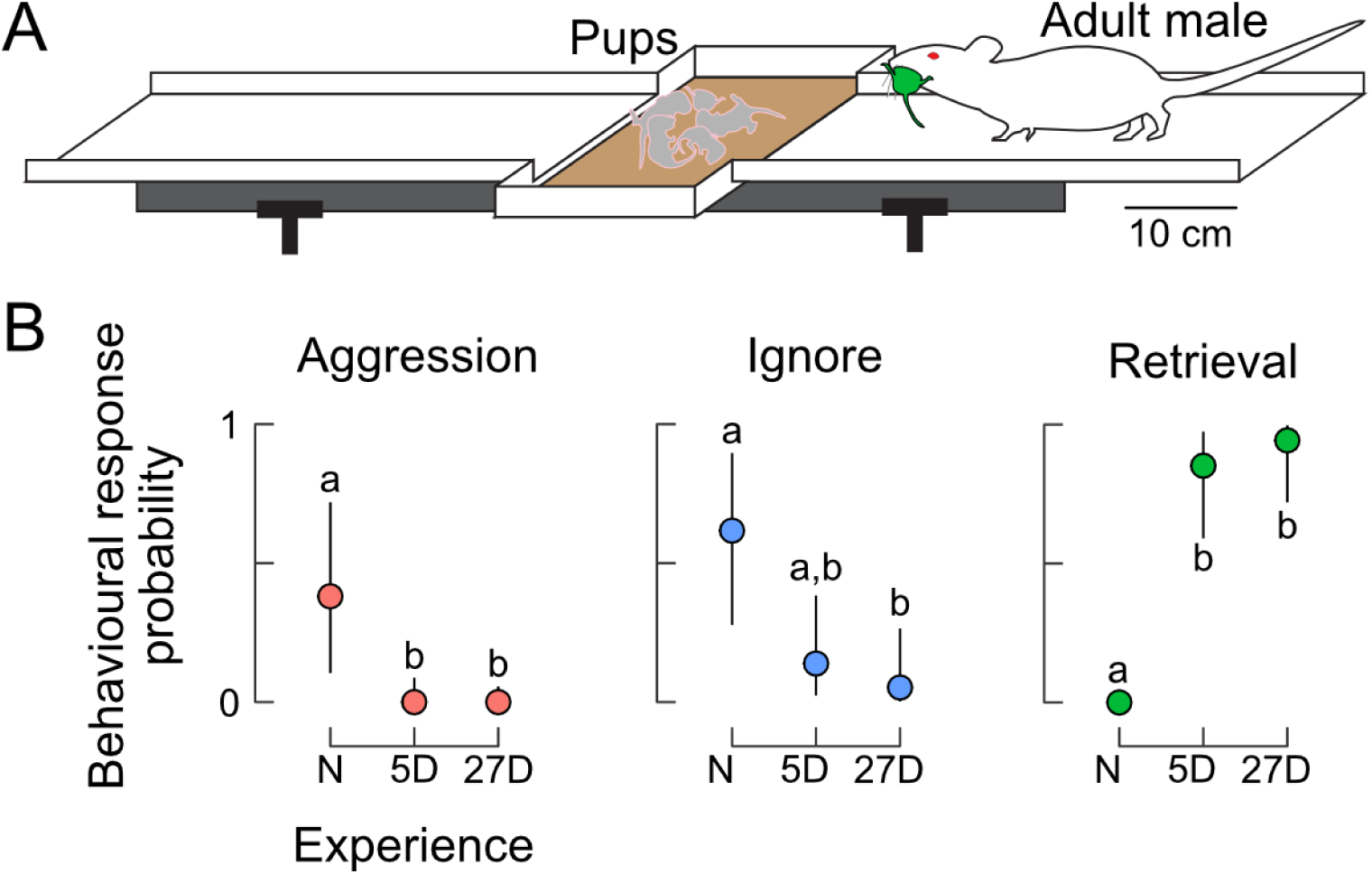
Behavioral responses towards pups. **A.** Experimental setting for the pup-retrieving tests on a running board with a central nest depression. Responses were scored as aggression (attacking pups), ignoring pups, or retrieval of pups (returning pups to the nest). **B.** Probabilities of behavioral expression in naïve (N) males and in fathers with five (5D) or 27 days of experience (27D). Aggression declined immediately from N to 5D fathers. Although fathers occasionally still ignored pups, the probability of retrieval was very high. Different letters indicate statistical differences between the groups of males. Sample size was n=8 for N, n=12 for 5D, and n=12 for 27D.

### Immunocytochemistry and microscopy

Immediately following the pup retrieval test, males were euthanized by cervical dislocation and their brains quickly dissected and frozen over liquid nitrogen. Brains were mounted using O.C.T compound and frontal sections were serially cut (30 μm) using a cryostat (Thermo Scientific Microm HM560), and immediately mounted on slides to keep track of the left and right sides of the brain. Sections were fixed in 4% paraformaldehyde in 0.1 M phosphate buffer (PB; pH = 7.4) for 60 min. Sections were rinsed three times for 5 min with 0.1 M PB in between all subsequent processing steps. Slides were incubated with 0.2% Triton X-100 (Sigma) in 0.1 M PB for 45 min to permeabilize cells and with 1% H_2_O_2_ (Merck) in 0.1 M PB for 20 min to block endogenous peroxidase. In a humidified chamber, the sections were treated with 2% normal goat serum in 0.1 M PB for 60 min to block non-specific staining, then with the polyclonal rabbit anti-aromatase antibody (Abcam Cat# ab35604, RRID:AB_867729; 0.002 mg/ml in 2% normal goat serum) at 4°C for 72h. Sections were then incubated with the secondary antibody from goat against rabbit horseradish peroxidase (Dako, Cat# PO448, 0.00125 mg/ml in 0.1 M phosphate buffer) at room temperature for 60 min. Immunoreactivity was revealed by incubating sections in 0.015% diaminobenzidine tetrachloride (DAB; Sigma), 0.023% NiCl, and 0.013% H_2_O_2_ in 0.1 M PB for 7 min. Sections were rinsed in PB, dehydrated in a series of alcohols and xylene, and coverslipped with Entellan (Merck). At the time of purchase, the primary antibody was recommended for aromatase immunohistochemistry on mouse tissue and the supplier provided positive quality controls with mouse ovarian tissue. Dutta et al. (2014) also performed a Western blot with mouse tissue that confirmed specificity of the antibody. Previous studies have used this antibody to detect aromatase immunoreactivity in mouse ovary (Laws et al., 2014), brain stem (Ji et al., 2017) and auditory cortex (Charitidi and Canlon, 2010). We tested the specificity of the secondary antibody labeling by incubating sections with PB instead of the primary antibody. In all these control sections, immunoreactivity was completely absent (Fig. 2A).

**Figure 2.**
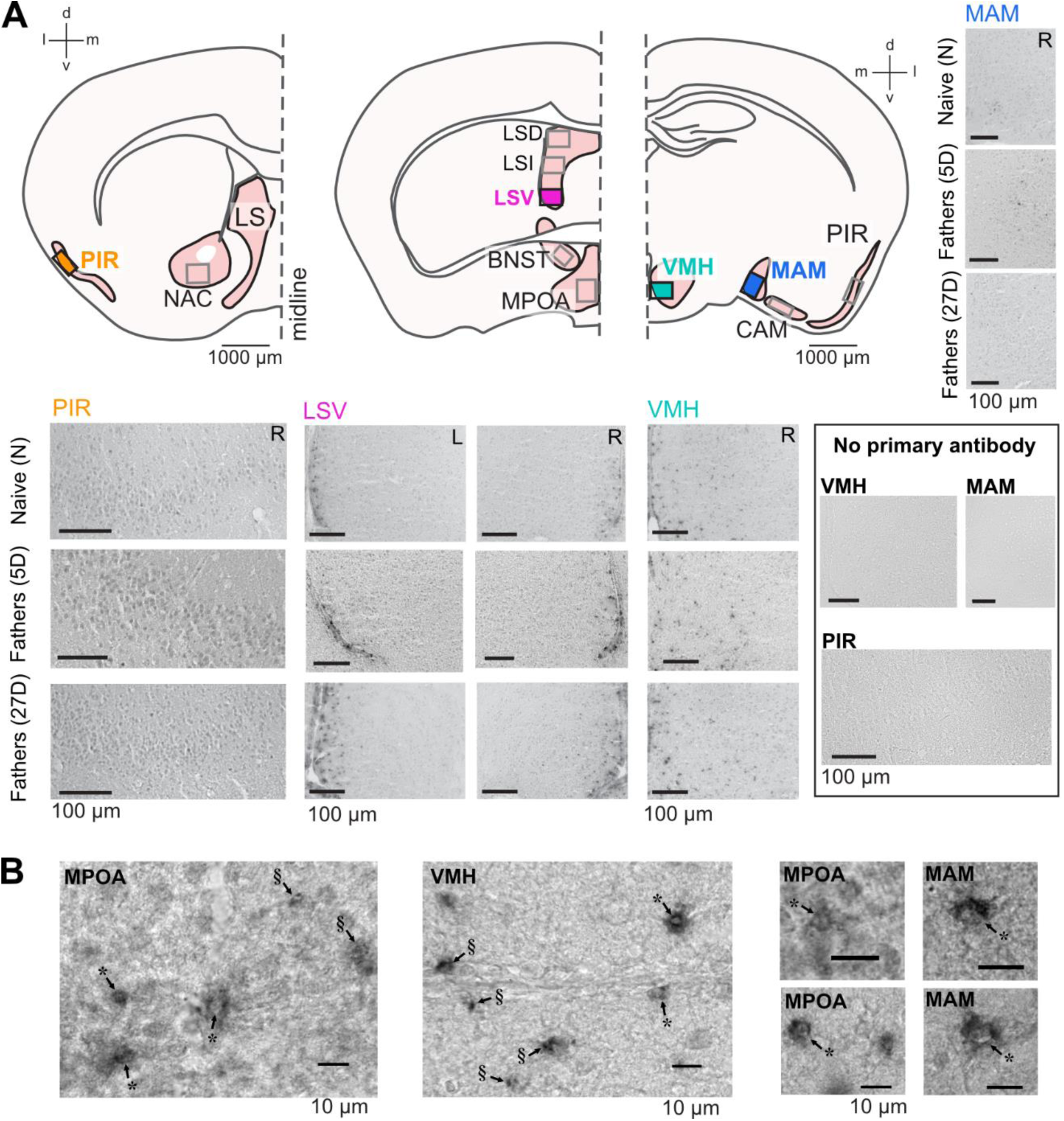
Aromatase immunolabeling in the paternal brain. Cross sections (**A**) of the brain areas analyzed here (Bregma coordinates: 0.98 mm; 0.14 mm; and −1.34 mm). Boxes indicate approximate areas that were counted. Representative images from areas highlighted in (**A**), showing aromatase immunostaining in naïve and experienced fathers (5D and 27D) in the left (L) and right (R) hemisphere and lack of immunostaining in the no primary antibody controls. High magnification (**B**) images of examples of aromatase expressing cells in the MPOA, VMH, and MAM showing cytoplasmic expression and black granulae. In cases in which the granulae were located perinuclear to an identifiable cell nucleus (marked with *) the cells were counted as aromatase-positive. In cases in which the granulae could not be associated with a cell nucleus (marked with §) the structure was not counted as an aromatase-positive cell.

Sections were independently analyzed by two blinded experimenters. In view of varying non-specific background labeling, the experimenters started the analysis by checking many sections from all animal groups in order to agree which kind of labeling would be considered aromatase-specific. Brain areas were identified with 10x objective magnification, aromatase-positive cells were identified by focusing with a 40x objective magnification. Cells were considered as expressing aromatase if they were (1) more intensely stained with shades of gray than the background, and (2) contained black granulae often grouped around the cell nucleus (Fig. 2B). In cases in which we observed a single or many black granulae without an associated cellular structure in shades of gray, the structure was not counted as an aromatase-positive cell (Fig. 2C). This procedure led to (a) a high concordance of counting results of the two experimenters, (b) counts of only definite aromatase-positive cells, and (c) relatively low numbers of aromatase-positive cells per evaluated brain area in a given section.

Sections were analyzed with a light microscope (Axiophot, Zeiss Germany). The brain areas in which aromatase-positive cells were counted are listed in Table 1. Areas were matched to the mouse brain atlas plates and associated Bregma coordinates (Paxinos and Franklin, 2001) and anatomically matched across all experimental animals. These areas largely overlap with those showing ERα positive cell dynamics during the acquisition of pup-caring experience in male (Ehret et al., 1993) and female (Ehret and Buckenmaier, 1994) mice.. Aromatase-positive cells were counted separately in the right and left hemispheres of the brain. Counts per section for a given animal, brain area and hemisphere were averaged between the maximum number of consecutive sections that could be matched across all animals, i.e., two sections for LS, BNST, VMH, and nucleus accumbens (NAC), or three sections for PIR, MPOA, cortical amygdala (CAM), and medial amygdala (MAM). These average counts from the individual animals entered further statistical analyses concerning the experimental groups of animals (N, 5D, 27D) and comparisons among the groups.

**Table 1.**
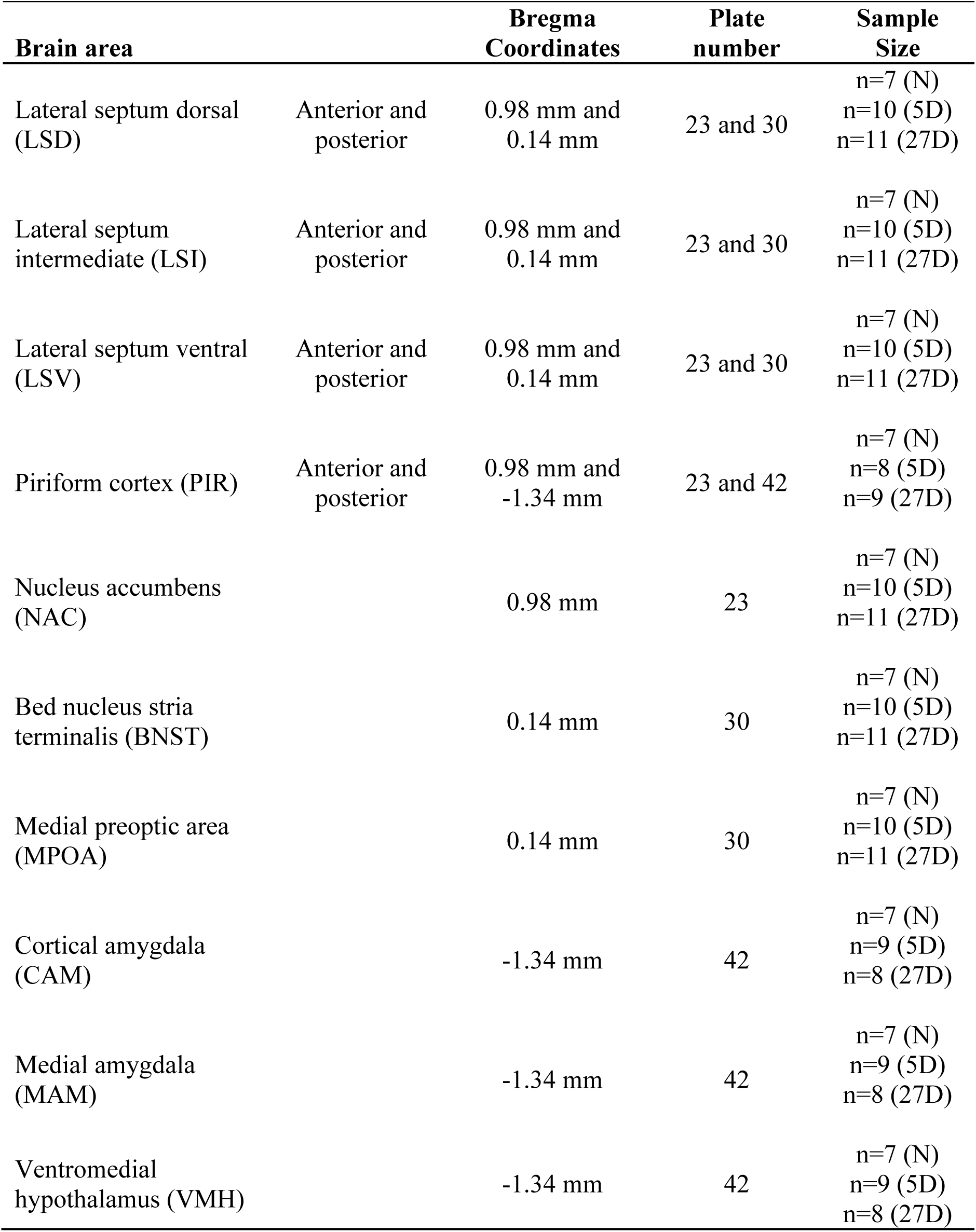
Brain areas analyzed for aromatase expression, their Bregma coordinates and plate numbers from the mouse brain atlas (Paxinos and Franklin, 2001), and sample sizes per group per brain area.

| Brain area |  | Bregma Coordinates | Plate number | Sample Size |
| --- | --- | --- | --- | --- |
| Lateral septum dorsal (LSD) | Anterior and posterior | 0.98 mm and 0.14 mm | 23 and 30 | n=7 (N)<br>n=10 (5D)<br>n=11 (27D) |
| Lateral septum intermediate (LSI) | Anterior and posterior | 0.98 mm and 0.14 mm | 23 and 30 | n=7 (N)<br>n=10 (5D)<br>n=11 (27D) |
| Lateral septum ventral (LSV) | Anterior and posterior | 0.98 mm and 0.14 mm | 23 and 30 | n=7 (N)<br>n=10 (5D)<br>n=11 (27D) |
| Piriform cortex (PIR) | Anterior and posterior | 0.98 mm and -1.34 mm | 23 and 42 | n=7 (N)<br>n=8 (5D)<br>n=9 (27D) |
| Nucleus accumbens (NAC) |  | 0.98 mm | 23 | n=7 (N)<br>n=10 (5D)<br>n=11 (27D) |
| Bed nucleus stria terminalis (BNST) |  | 0.14 mm | 30 | n=7 (N)<br>n=10 (5D)<br>n=11 (27D) |
| Medial preoptic area (MPOA) |  | 0.14 mm | 30 | n=7 (N)<br>n=10 (5D)<br>n=11 (27D) |
| Cortical amygdala (CAM) |  | -1.34 mm | 42 | n=7 (N)<br>n=9 (5D)<br>n=8 (27D) |
| Medial amygdala (MAM) |  | -1.34 mm | 42 | n=7 (N)<br>n=9 (5D)<br>n=8 (27D) |
| Ventromedial hypothalamus (VMH) |  | -1.34 mm | 42 | n=7 (N)<br>n=9 (5D)<br>n=8 (27D) |

### Statistical analyses

All statistical analyses were carried out using R v4.2.0 (R Core Team, 2022). **Behavior:** Male behavior in response to pup presentation was classified as ignoring, attacking, or retrieving a pup. The outcomes of behavioral trials were analysed as the multinomial probability of exhibiting a given response to pup presentation (aggression, ignore, retrieval) as a function of paternal experience (experience as ordered factor, N < 5D < 27D). To perform pairwise comparisons between the groups for each behavior, we calculated the probability that the difference in the posterior predictive intervals between groups was significantly directional. We then corrected for multiple comparisons using the false discovery rate.

#### Aromatase immunolabeling

Aromatase cell counts were conducted in two data sets, which were combined to increase statistical power for comparisons. The data sets were obtained with the same immunocytochemical protocols and the same chemicals, however, the experiments were performed as two separate cohorts. We observed some count variation between the data of the cohorts (e.g., for the PIR −1.34 data; see Supplemental Fig. 2), and so each data set was standardized by the mean and variance of the paternal groups (5D and 27D; the groups for which the datasets were overlapping). Sections for some brain areas were missing in some mice. The pattern of missingness would have led to omitting eight of 56 individuals (14%). In order not to totally exclude these animals from the data analysis, we elected to impute the missing data (16 cases, 5.7% of 280 total region counts) using predictive mean matching (van Buuren and Groothuis-Oudshoorn, 2011; R package mice). We generated five such candidate data sets and integrated over the resulting uncertainty.

Our statistical model of aromatase variation was designed to address non-independence arising from correlated variation within individuals among areas. Such non-independence could be an especially important factor when comparing identifiable subregions of an area, such as in the piriform cortex and each part of the lateral septum. We therefore examined variation in normalized cell counts among all brain areas simultaneously (multivariate ANOVA), treating individual mouse identification (ID) as a correlated group (random) effect to account for hemisphere differences within individual. Responses were modelled as *t-*distributed to increase robustness against outlier counts. The interaction of Experience x Hemisphere was examined together.

All models were fit using the R package *brms* (Bürkner, 2017) using four chains of 2500 iterations each, and a thinning rate of 5. We assumed uniform priors throughout. Model convergence was assessed by 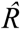 = 1 (Vehtari et al., 2021); the 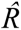 statistic was computed within each imputed data set. We calculated the posterior means of each data set based on the fitted statistical model (*emmeans* package; Lenth, 2022), and used these to examine the posterior distributions of contrasts between hemispheres (left *minus* right for each brain area and experimental group of animals) and between the experimental groups for each brain area and hemisphere, i.e. 5D minus N (5D-N), 27D minus N (27D-N), 27D minus 5D (27D-5D). These data are plotted in Figure 3. Significance of each contrast was judged as a Bayesian probability of direction P > 0.975 (Makowski et al., 2019), which is an index of effect existence, i.e., our level of certainty in whether a reported change is different from zero, either positive or negative. This threshold closely corresponds to the two-tailed p-value of 0.05, and so for simplicity, we report the p-value. Plots were generated using the *ggplot2* package (Wickham, 2016).

**Figure 3.**
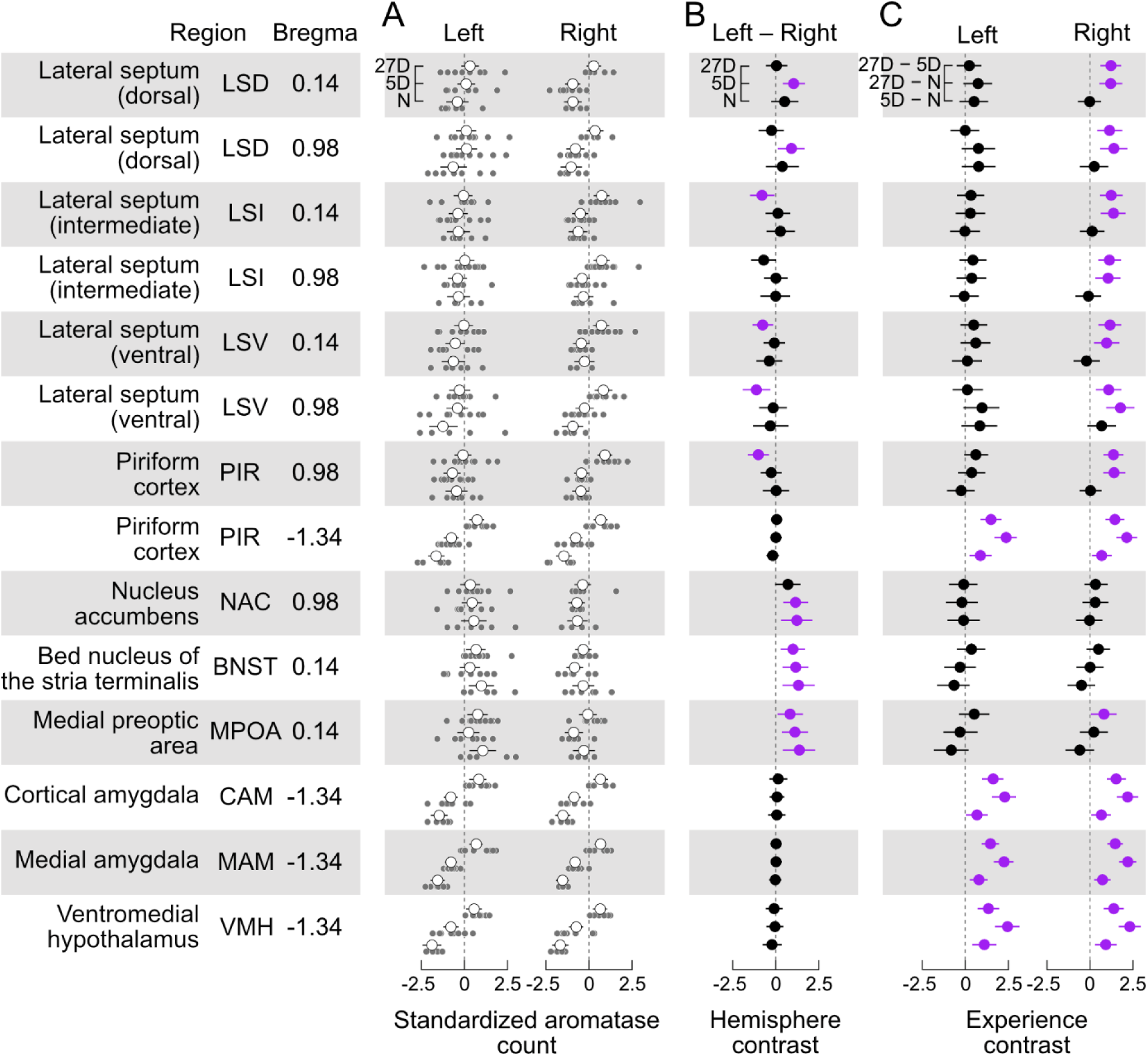
Aromatase expression differentially depends on paternal experience and brain hemisphere. **A.** Standardized cell counts (light grey circles) and means with ± 95% confidence interval (white circles) for each investigated brain area separately for the left and right sides of the brain and levels of pup-experience (N, 5D, 27D). **B.** Hemisphere contrast (left – right, with 95% confidence intervals) of the standardized counts of aromatase positive cells for each brain area and experimental group. Contrasts with intervals significantly different (p < 0.05) from zero are highlighted in purple (see text for description). We found evidence for left-hemisphere dominant lateralization (left minus right difference was positive) of aromatase cell counts both independent of (BNST, MPOA, and most likely NAC) and dependent on pup-experience (dorsal LS), and an experience-dependent right-hemisphere dominant lateralization in the intermediate and ventral LS and anterior PIR (Bregma 0.98). **C.** Experience contrast (5D–N; 27D–N, 27D–5D, with 95% confidence intervals) for the standardized counts of aromatase positive cells, separately for each hemisphere and brain nucleus/area. We found that the occurrence of aromatase positive cells in posterior (Bregma - 1.34) PIR, CAM, MAM, and VMH in both brain hemispheres significantly increased with the experience level. Expression of aromatase in the LS and anterior PIR significantly increased only in the right hemisphere of the 27D group. No changes were observed in the NAC and BNST. Contrasts with intervals significantly different (p < 0.05) from zero are highlighted in purple (see text for description). Sample sizes are given in Table 1.

Residual correlations among areas represent relationships that are not explained by the other population- and group-effects, including correlated expression and unexpected technical covariation. Inspection of residual correlations can therefore inform biological hypotheses. However, direct interpretation of the residual correlation matrix is challenging because the correlation between two brain areas could be due to mutual covariation with another area. We dichotomized relationships among brain areas by studying the matrix of *partial* correlations (de la Fuente et al., 2004; Shipley, 2000). We constructed an initial graph in which edges were created between all nodes with significant (p < 0.05) pairwise correlations. Edges were iteratively pruned by testing whether the association remained statistically significant after testing it against all other paths between two variables, up to third order. The independence test was determined from the posterior distribution of residual correlation matrices, and implemented in the R package pcalg (Kalisch et al., 2012). The result is an undirected, acyclic graph representing a network of associations among brain areas. The graph is shown in Figure 4. The large number of tests may be conservative and may reject true associations, so we also report tentative links observed when setting a less stringent threshold of p < 0.10.

**Figure 4.**
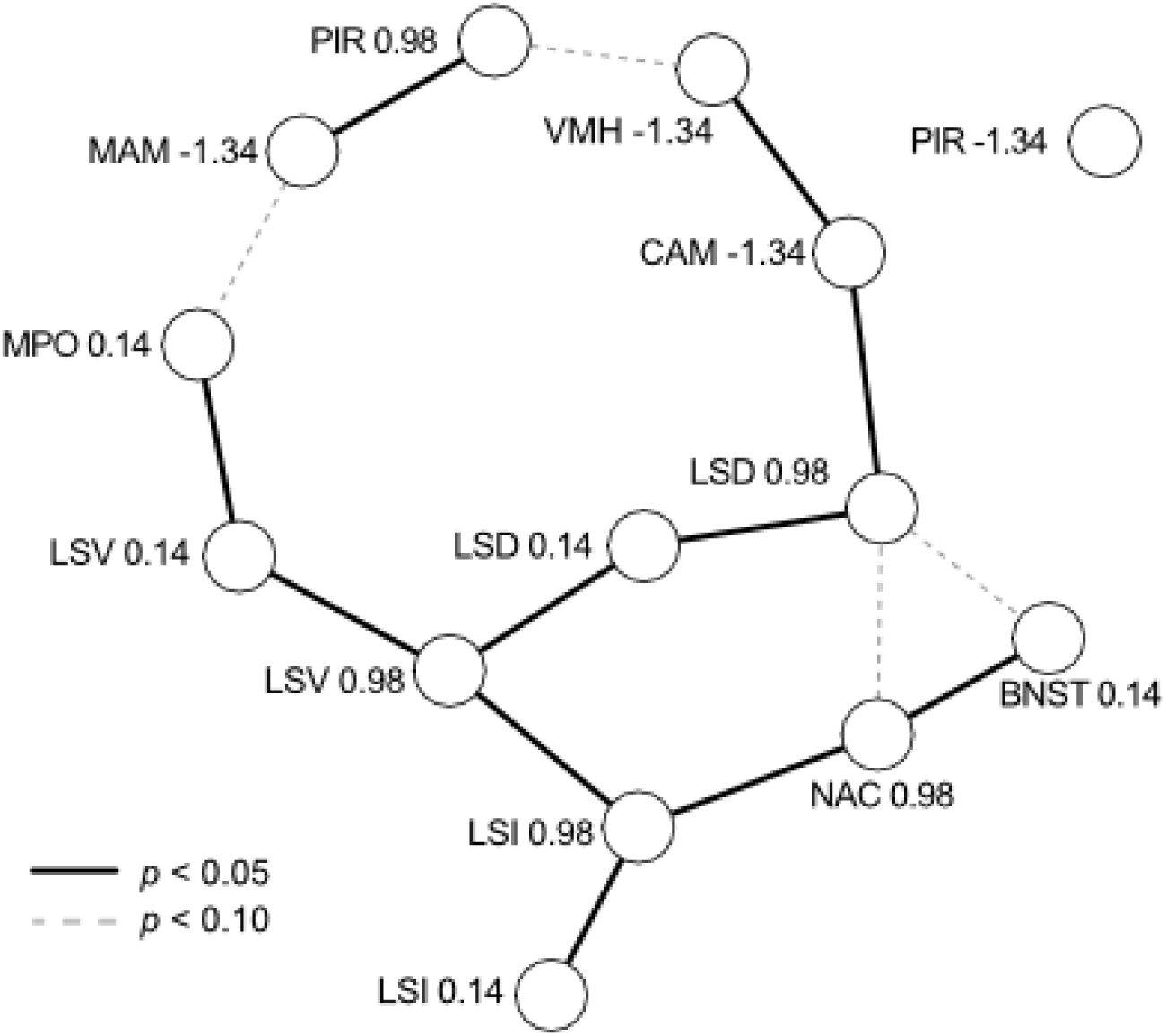
Residual correlations among brain nuclei/areas viewed as a network. Each nucleus/area at each Bregma coordinate (see Table 1) is represented by a node, with edges representing significant partial correlations (solid for p < 0.05, dashed for p < 0.10). Residual correlations were positive between all brain areas, with the sole exception of CAM (−1.34) and LSD (0.98), which were negatively correlated. PIR (−1.34) was not correlated with any other brain area.

#### Pup ignoring and aromatase immunolabeling

Pup retrieval and aggression behaviors are completely colinear with the treatment groups (see below), so we could not examine behavioral differences simultaneously with experience. Pup ignoring, however, was observed in all experimental groups of mice. Therefore, we sought to find factors predicting why some males, regardless of experience, ignore pups (or why some males react to pups). Behavior toward pups was coded as reacting (aggressiveness or retrieval) or ignoring, and we then conducted a similar analysis to that described above for aromatase cell counts. All males were examined together without considering differences in experiences or hemispherical variation in aromatase cell counts.

## Results

### Male mice retrieve pups after acquiring parental experience

Only males with paternal experience retrieved pups (Fig. 1A). Figure 1B shows that naïve males with no previous sexual experience and pup contact mostly ignored the pups (n = 5/8) or were aggressive (n = 3/8). Most fathers with 5 days (5D) or 26-27 days (27D) of pup-experience retrieved pups (n = 10/12 and 11/12, respectively) and only few ignored pups (n = 2/12 and 1/12, respectively). No father was aggressive.

The probability of observing aggressive behavior toward pups was significantly higher in naïve males compared to fathers regardless of experience (5D and 27D, Fig. 1B). The probability of other non-paternal behavior like ignoring pups was higher in naïve males compared to 27D fathers. Parental behavior measured as pup retrieval was significantly higher in fathers (5D and 27D) compared to naïve males. There were no significant differences in the probability of pup-retrieving between the groups of fathers, 5D and 27D (Fig. 1B).

### Aromatase expression in the male mouse brain, general picture

We found aromatase-positive cells in each examined area and with each level of experience. In absolute terms, largest numbers of aromatase positive cells per section were seen in PIR, LSI, MAM, and CAM (see unstandardized data in Supplement, Fig. 2). Examples of aromatase labeling in the LSV, PIR, VMH and MAM of experimental groups N and 27D showed qualitative differences between the groups and between left and right sides of the brain (Fig. 2). Standardized cell counts together with the group means are shown in Fig. 3A along with our estimates of the posterior differences in aromatase counts by hemisphere (Left – Right, Fig. 3B), and between levels of experience (Fig. 3C).

### Lateralized occurrence of aromatase positive cells

We found evidence for significantly greater numbers of aromatase positive cells in the left hemisphere of the MPOA and BNST in all three experimental groups (Left – Right > 0, Fig. 3B). The NAC exhibited a similar trend to MPOA and BNST, but with a just insignificant contrast in the 27D group. We did not detect significant lateralization of aromatase counts in any of the experimental groups in the amygdala (CAM and MAM), VMH, or posterior PIR (Bregma −1.34). In the other brain areas, the appearance of lateralization strongly depended on the level of pup-experience. Long-term experience (27D) led to greater numbers of aromatase positive cells in the right hemispheres of anterior PIR (Bregma 0.98), the LSV, and the anterior LSI (Bregma 0.14). Posterior LSI (Bregma 0.98) exhibited a similar trend that was just non-significant. In the LSD, pup-experience had a short-term effect (5D) during which counts in the left hemisphere were greater. This effect was not present in the 27D group (Fig. 3B).

### Experience-dependent expression of aromatase

In all cases in which we found significant changes of aromatase counts with pup-experience, the number of aromatase positive cells increased (Fig. 3C). This experience-dependent increase was pervasive for the amygdala (CAM and MAM), the VMH, and the posterior PIR (Bregma −1.34). For these areas, the increase in aromatase positive cells was present in both hemispheres and with increasing experience, i.e., experience contrasts had significant positive values for both brain hemispheres and all experimental groups. In the LSD, LSI, and LSV (both Bregma coordinates), and the anterior PIR (Bregma 0.98), aromatase positive cells increased significantly only in the right hemisphere of 27D males. Experience-dependent effects were absent in the NAC, the BNST, and the MPOA, except for a significant effect in the right MPOA of 27D males, i.e., 27D males had significantly more aromatase positive cells than 5D males (Fig. 3C).

### Graphical analysis of covariation among brain areas

We examined the unexplained (residual) variation in aromatase cell numbers among brain areas as graphically shown in Figure 4. We found positive residual correlations between all brain areas, with the sole exception of CAM (−1.34) and LSD (0.98), which were negatively correlated. Three subgroups of covariation are obvious. First, the LS is highly connected and each subregion (D, I, V) is correlated across Bregma coordinates and to at least one other subregion. The LS appears to form the backbone of variation, to which other nodes are connected. This includes subgroups comprised of MPOA (connected to LSV), of NAC and BNST (connected to LSI), and of CAM and VMH (connected to LSD). Second, the subgroup of MAM and anterior PIR appears to be weakly connected to other parts of the graph (MPOA and VMH), because the edges are significant only with p < 0.10. Third, posterior PIR did not appear to be even weakly correlated with any other analyzed brain area despite the increase in aromatase expression with experience that was similar to CAM, MAM, and VMH (see Fig. 3).

### Variation in aromatase expression in males that react or ignore pups

The loss of aggressiveness towards pups and the gain of pup retrieval behavior starkly marks the transition to fatherhood (Fig. 1). Nonetheless, some males appear to ignore pups, and even one second-time father failed to retrieve pups during the pup retrieval test. We therefore examined if variation in aromatase cell counts in any area might predict whether a male reacted to or did not react to pups (contrast of reacting – ignoring mean cell counts, Fig. 5). Reacting males had significantly higher numbers of aromatase cells in the MAM (p=0.011), and a weak tendency to higher numbers in the CAM (p=0.06).

**Figure 5.**
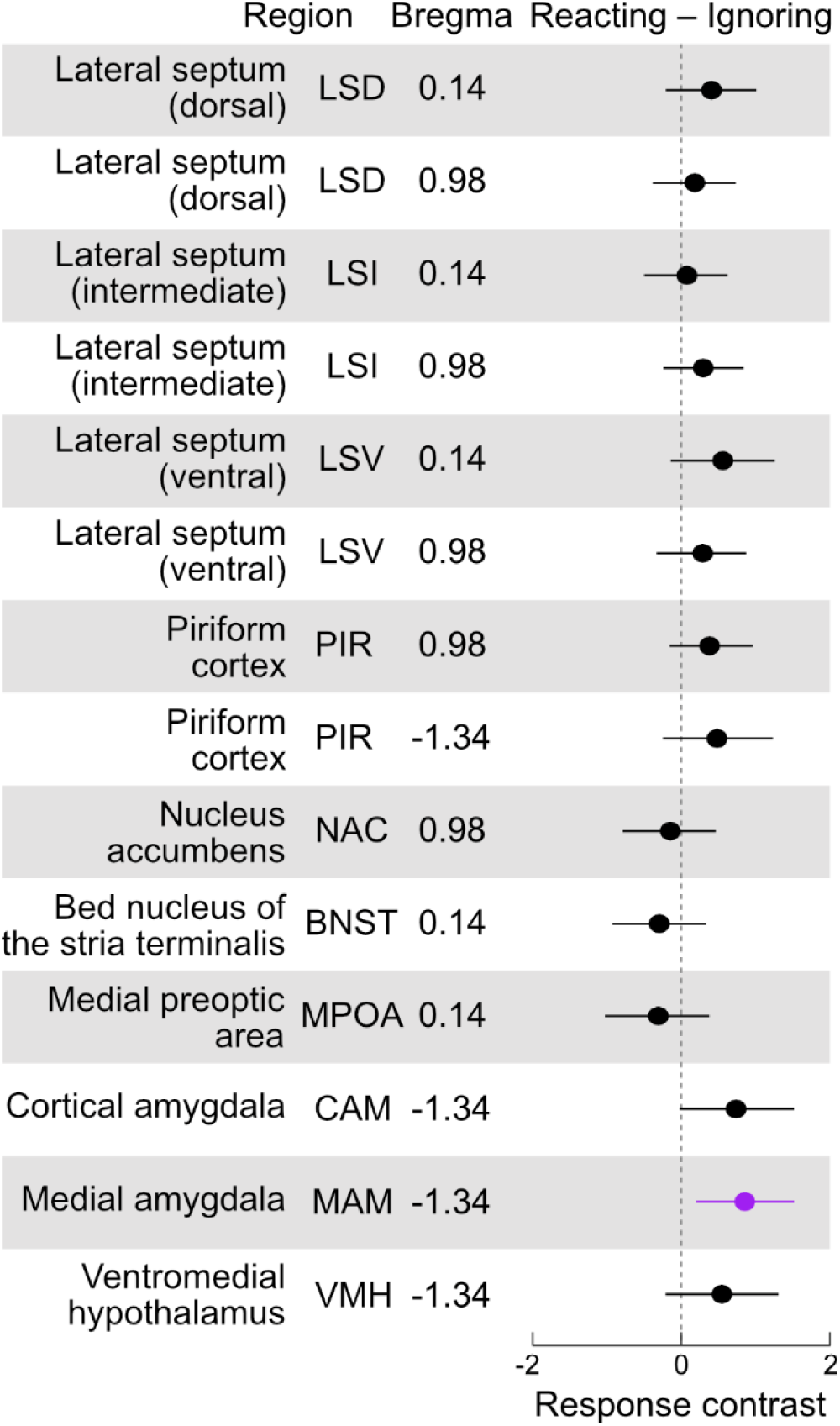
Aromatase expression in males that react to pups compared to those ignoring pups. Contrasts (reacting – ignoring, with 95% confidence intervals) of the standardized counts of aromatase positive cells for each brain area. Contrasts with intervals significantly different (p < 0.05) from zero are highlighted in purple (see text for description). We found that reacting males had significantly higher numbers of aromatase cells in the MAM (p=0.011), and a weak tendency to higher numbers in the CAM (p=0.06).

## Discussion

In the present study, we found that aromatase, which catalyses the conversion of testosterone to estrogen, is expressed throughout areas of the limbic system of male mice, regardless of their experience with pups. However, we identified important factors contributing to hierarchical variation in expression levels among individuals. First, the dramatic changes in behavior associated with paternal experience were associated with changes in aromatase expression in areas/nuclei of the limbic system. However, not only were these changes area-specific, but also hemisphere-specific (summarised in Fig. 6). Finally, layered on these widespread patterns, we detected correlated aromatase levels among brain areas within individual mice. We briefly review the role of aromatase and the limbic areas to discuss how such expression patterns may influence, and be regulated by, paternal behaviors and experience.

**Figure 6.**
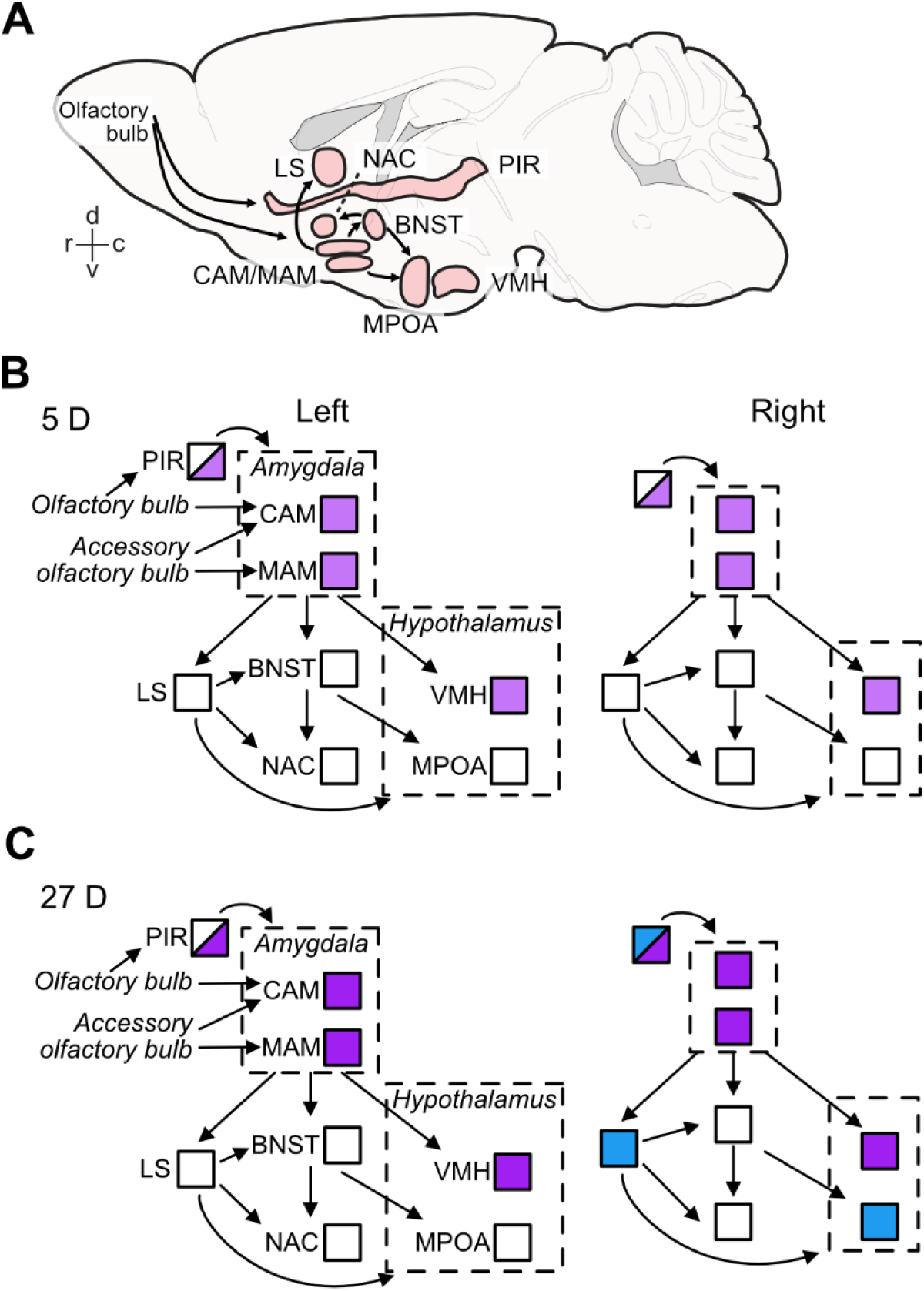
Effects of paternal experience on aromatase expression in areas of the limbic system. Sagittal view of the brain areas analyzed here (A) and summary of the changes in aromatase expression in 5D **(B)** and 27D **(C)** males in the left and right hemispheres of the brain nuclei/areas analyzed in the current study. **A.** Positions and extensions of nuclei/areas are illustrative only and do not indicate anatomical relationships. The connectivity denoted in (A) shown by the arrows are exemplars for main information pathways (Numan, 2020; Kohl et al., 2017). **B.** Pup experience in first-litter fathers (5D group) increased the numbers of aromatase positive cells (indicated by light purple) relative to naïve males in the amygdala (CAM and MAM), hypothalamus (VMH), and in posterior piriform cortex (PIR, half shading indicates these partial changes). **C.** Second-litter fathers (27D group) had further increased aromatase expression in the same areas as 5D males (indicated as darker purple), i.e., the amygdala (CAM and MAM), hypothalamus (VMH), and posterior piriform cortex (PIR) in both sides of the brain. In the right hemisphere, the number of aromatase positive cells increased (indicated by blue) in the anterior PIR, lateral septum (LS) relative to naïve and 5D males, and potentially in the MPOA relative to 5D males.

### Aromatase and the expression of paternal care

Aromatase expression in the male mouse brain has been shown via immunostaining with several protocols using different antibodies (Akther et al., 2015; Foidart et al., 1995), partly with enhancement through transgenic techniques (Stanić et al., 2014). In agreement with the published data, we found aromatase expression in all studied brain areas of our three experimental groups of male mice (Fig. 3). Differences in the levels of aromatase expression between studies relate to differences in the staining methods, data analysis, and the social environment of the animals prior to and during the behavioral tests. For example, in our study, dynamics of paternal behavior and associated changes in aromatase levels have been examined while the fathers were continually housed with their female mate except during the pup-retrieval tests. This is different from the study of Akther et al. (2015) in which the relationship between pup-retrieving behavior in males and the dynamics of aromatase levels was tested throughout different ethological contexts (presence, absence and reintroduction of their female mates and pups). In that previous study, brain aromatase expression was highest when the fathers were housed with their female partner, but without their pups, followed by fathers housed with the whole family (Akther et al., 2015). Therefore, the observed changes in aromatase levels in that study can be related to male-female interaction. Nevertheless, the strong reduction of pup-retrieval in fathers after injection with an aromatase inhibitor (letrozole; Akther et al., 2015) shows, along with the results of our present study, that proper levels of aromatase expression are integral to the display of paternal behavior in mice.

Interestingly, circulating testosterone is negatively correlated with paternal behavior in various rodent species (reviewed in Bales and Saltzman, 2016; Numan and Insel, 2003) and testosterone can inhibit paternal care in mice (Okabe et al., 2013). Our study did not examine the source of testosterone, the substrate of aromatase in the brain. However, previous studies have shown that changes in circulating testosterone may not be reflected locally in the brain. Neurosteroids can be locally produced, and levels in the brain may be independent of circulation or gonadal status (Giatti et al., 2019; Hojo et al., 2008, 2004).

In the following, we discuss the observed presence and changes of aromatase levels in the studied brain areas of the limbic system in order to see how these areas may contribute to changing male behavior towards pups from aggressive or ignoring in naïve males to caring in experienced fathers (Fig. 1). And further, the comparison of dynamics of aromatase-positive cells with the dynamics of ERα-positive cells in the given brain areas will show in how far they are in parallel and support the hypothesis about aromatase influencing indirectly (via estrogen synthesis and ER binding) paternal behavior.

### Amygdala and ventromedial hypothalamus

Three nuclei, the MAM, CAM, and VMH were most affected by pup experience with increases in the number of aromatase positive cells from N to 5D to 27D animals (Fig. 3 A and C; Supplement Fig. 2). These areas are associated with the regulation of aggression compared to sexual and parental behavior (e.g., Chen et al., 2019; Dwyer et al., 2022; Hashikawa et al., 2016; Lee et al., 2014; Sheehan et al., 2001; Unger et al., 2015; Yang et al., 2017). Important changes in aggression in the course of becoming a caring father are mediated by olfactory signals from the vomeronasal organ via the accessory olfactory bulb to the CAM and MAM (e.g., Hashikawa et al., 2016) and concern the switch from aggressive to caring behavior towards pups (Tashikawa et al., 2013). Concomitantly, aggressive motivation in fathers increases against male intruders in order to protect their own pups from potential pup killers as shown in house mice (Shabalova et al., 2020), Californian mice (Trainor et al., 2008) and prairie voles (Getz et al., 1981). This increased aggression towards intruders was not mediated by changes in ERα or ERß expression in limbic centers, but reflected in increased neuronal activation (FOS-expression) in the MPOA, MAM and, potentially the VMH in California mice (Trainor et al., 2008). In house mice fathers, aggressive behavior toward intruders is regulated by oxytocin and activation of neurons in areas of the hypothalamus (periventricular and supraoptic nuclei; Shabalova et al., 2020). Future work could examine whether changes in aggression in naïve vs fathers are also related to aromatase levels in the amygdala and VMH.

In contrast to aromatase expression (present study), the number of estrogen receptors (ERα) in MAM, CAM, and VMH did not change with parental experience in male mice (5D or 27D; (Ehret et al., 1993). Together, this indicates that the increased presence of local estrogen synthesis does not facilitate the expression of ERα, as has been shown in the VMH (Devidze et al., 2005; Malikov and Madeira, 2013), especially in combination with parental experience (Koch, 1990). Very limited work has been done on how other estrogen receptors are involved in paternal care in rodents (Hyer et al., 2017; Torres et al., 2024). It is possible that other subtypes (ERβ and membrane receptor GPER) are involved in the expression of parental behavior. Our work indicates that parental experience is associated with changes in aromatase expression, and this can potentially affect the balance of androgen to estrogen levels in the brain. Future work should also examine whether androgen receptors, expressed in the same brain areas as aromatase (Dart et al., 2024), are implicated in parental behavior (e.g., see work in Mongolian gerbils; Martinez et al., 2019). We suggest that the increase in aromatase expression in the MAM, CAM, and VMH in new (5D) and highly experienced (27D) fathers (Fig. 6) supports the control of instinctive behavior (Honda et al., 2011) including inhibition of infanticide, via increased local synthesis of estrogens on neurons (e.g., Cornil et al., 2006, 2013).

### Piriform cortex

In house mouse mothers, the performance of pup-retrieving does not depend on olfactory signals from the main olfactory bulb and its projections to the PIR (Koch and Ehret, 1991). Pup-naïve females, however, needed this olfactory information for learning to become as maternal as the mothers (Ehret and Buckenmaier, 1994; Koch and Ehret, 1991). While learning pup cues for becoming maternal, virgin females with maternal experience have an increase in ERα positive cells in the posterior PIR compared to females without experience (also in the entorhinal cortex; Koch, 1990). Similarly, as males became caring fathers, ERα positive cells become visible in the posterior PIR (also in the entorhinal cortex; Ehret et al., 1993). In view of the significant increase of aromatase expression in the posterior PIR with increasing paternal experience (Fig. 3C), it is possible that in PIR, unlike MAM, CAM, and VMH, estrogen may have induced its own receptors (tested for ERα) and enhanced olfactory learning of pup cues for increased paternal performance.

The significant increase of aromatase positive cells only in the right hemisphere in the anterior PIR of 27D fathers does not directly correlate with any evidence known to us. In studies on ERα expression in the limbic areas of female and male mice under various conditions of the reproductive cycle, experience with pups, and intact or lesioned olfaction, no hemisphere differences have been reported in any area (Ehret et al., 1993; Ehret and Buckenmaier, 1994; Koch, 1990; Koch and Ehret, 1991). The splitting of PIR in regions with experience-dependent lateralized (anterior PIR) and non-lateralized (posterior PIR) aromatase change shows compartmentalization of the PIR. Functional relationships need to be elucidated in further studies.

### Lateral septum

The main effects of pup-experience on aromatase expression in the LS were changes in the left-right balance (Fig. 3B) and significant increase of aromatase-positive cells only in the right LS of highly experienced (27D) fathers (Fig. 3C). As in the PIR, ERα positive cells in the LS become visible in caring fathers compared to pup-naïve males (Ehret et al., 1993). Since hemisphere differences in ERα positive cells have not been found, we cannot directly relate increased right-hemisphere aromatase expression with non-lateralized increase of ERα expression in the LS of both hemispheres. Lateralized input to the rostral parts of the LSD and LSI comes from the hippocampal CA1 and subiculum region (work in mice and rats; Rizzi-Wise and Wang, 2021; Risold and Swanson, 1997). In rodents, the CA1 shows lateralization in synapse morphology and receptor distribution (Shinohara et al., 2008; Xiao and Jordan, 2002) and left-right dissociation of memory processes (Shipton et al., 2014) with right-hemisphere dominance of spatial memory in mice (Shinohara et al., 2012). Also, androgen receptor levels are higher in the left hippocampus in male rats (Xiao and Jordan, 2002). It is unclear, however, whether and how lateralized hippocampal input to the LS could have caused lateralized aromatase expression in the LS of 27D fathers (Fig. 3C). Right hemisphere lateralization is associated with the regulation of mood, affect, and stress, and with sustained and global arousal and attention (e.g., Ehret, 2006; Palomero-Gallagher and Amunts, 2022). Speculatively, our observed right-hemisphere LS bias in 27D fathers could be related to the stress of continuous arousal and steady involvement in a routine of pup-care over several weeks.

### Medial preoptic area, bed nucleus stria terminalis, and nucleus accumbens

The MPOA together with the BNST and their output via the NAC are essential for controlling parental behavior in rodents (e.g., Numan, 2006; Bridges, 2015; Wu et al., 2014). Significant increases have been observed in the MPOA both of the number of and area occupied by ERα positive cells in 5D and 27D pup-experienced father mice (Ehret et al., 1993) and, although absent in pup-naïve males, ERα-positive cells are present in the BNST of 27D fathers (Ehret et al., 1993; Koch and Ehret, 1989). In females, the number of ERα positive cells are also higher in the MPOA and BNST of one-day pup-experienced virgin females (Ehret and Buckenmaier, 1994). In the current study, we did not observe changes in the number of aromatase positive cells in the BNST and NAC and found only a small increase in the MPOA in just the 27D males relative to 5D. This suggests that changes in sensory processing and behavior with experience in MPOA, BNST, and NAC may be more dependent on regulation of estrogen signaling through binding to ERα than on local estrogen action after synthesis via aromatase. However further work on other estrogen receptors is needed. Brain areas that regulate social behavior, including parental care, are conserved in vertebrates (O’Connell and Hofmann, 2012). However, neuroendocrine ligand spatial profiles are more variable than receptor distributions within the social decision-making network among vertebrates, although there are regional differences (O’Connell and Hofmann, 2012). Our work supports experience-dependent regional differences and specifically that influences of MPOA, BNST, and NAC on parental behavior may depend on changes in ERα distribution, whereas the influences of VMH and amygdala may depend on changes in aromatase expression. Overall, our results point to an area-specific regulation of receptor and ligand synthesis with parental experience.

The brain areas MPOA, BNST, and NAC exhibited substantial lateralization of presence of aromatase-positive cells, biased toward the left hemisphere (Fig. 3B). Left-hemisphere dominance has been noted in a number of contexts, including maternal perception and processing of pup-ultrasounds, in the latter case, through cFos expression (Ehret, 1987; Geissler et al., 2016). Maternal pup retrieval behavior likewise is associated with left-but not right-hemisphere oxytocin receptor expression and action (Marlin et al., 2015). To what extent can these sources of left-hemisphere bias in females (primarily auditory cortex) inform our findings? The left-hemisphere bias in females has primarily been observed in the auditory cortex (Geissler et al., 2016; Marlin et al., 2015). Lateralization of cFos expression has not been observed in the neural responses to pup presence in the limbic system of female mice, MPOA and BNST included (Geissler et al., 2013). However, neural activation characterized by cFos labeling concerns the input side of neurons (e.g. Sheng et al., 1990, 1993). Provided such absence of hemisphere differences in activation at the input side of the MPOA and BNST also existed in males/fathers, we can hypothesize that the greater left hemisphere aromatase expression in the MPOA, BNST, may lead to a left-hemisphere bias of activation at the output of the MPOA and BNST after local estrogen action (enabled via aromatase) had taken place. Whether such a functional bias actually exists and how it may contribute to a preferred processing of pup ultrasounds in order to activate appetitive pup-searching behavior and possible left-hemisphere dominant ultrasound perception in pup-experienced male mice similar to mothers remains to be shown.

### Correlated individual variation in aromatase-positive cells between brain areas

The brain’s connectivity suggests that counts of active cells (e.g., c-Fos as indicative of brain activation) within individuals may be correlated across regions. For example, aromatase could be uniformly lower or higher in some individuals compared to others, or could vary highly non-uniformly across brain areas. We expect that such variation will contribute to individual differences in behavior, such as some fathers continuing to ignore pups (Fig. 1B). Our study found significant residual correlated variation among brain areas (Fig. 4), generally not depicted in the data shown in Fig. 3. Our analysis points to a central role of the LS in the variation of aromatase expression in the mouse brain. The LS has a central functional role in the brain integrating cognitive and emotional information in the regulation of social relationships and instincts (Besnard and Leroy, 2022; Menon et al., 2022; Rizzi-Wise and Wang, 2021) including parental behavior (Fleischer and Slotnick, 1978; Slotnick and Nigrosh, 1975). Interestingly, these partial correlations (shown in Fig. 4) reflect main neural connections of the subregions LSD and LSV. For example, there is input from the amygdala to the LS supporting the LSD-CAM partial correlation and there is an output from the LS to the MPOA supporting the LSV-MPOA partial correlation (Besnard and Leroy, 2022; Menon et al., 2022; Rizzi-Wise and Wang, 2021; Risold and Swanson, 1997). Thus, residual variation in aromatase expression (Fig. 4) appears to mirror functional connections and relationships of the LS. It is noteworthy that the posterior PIR and, to a lesser degree, the anterior PIR and the MAM remained outside this connectivity. Indeed, Supplementary Fig. 2 shows that posterior PIR (Bregma −1.34) was the sole case in which we observed substantial divergence in overall cell counts between the two cohorts of animals. Together, these observations on correlated individual variations indicate both coherent relationships between brain areas in a functional network and the presence of possibly unknown additional factors intrinsic to given experiments. In our present case, such factors modulated the numbers of aromatase-positive cells in PIR and MAM. Therefore, our study shows the usefulness of correlative modelling to assess and estimate the relationships between contributions of the members of a neuronal network to its overall functional properties. This is another example that correlative modelling supports the adequate interpretation of absolute data.

### Amygdala aromatase expression is related to reacting to pups

Our study design cannot fully link changes in individual behavior to changes in aromatase in the brain because the switch in the expression of both is so stark. We therefore sought to examine why some fathers *do not* appear to react to their own pups (i.e., they ignore their pups) and, like most naïve males, continue to ignore them. Although tentative, our results (Fig. 5) point to the possibility that the amygdala (especially the MAM) mediates strong emotional responses to pups and the particular expression of is dictated by the network of brain areas. It would be interesting to study the longitudinal responses of males (naïve and fathers) to pups, to address the individual repeatability of their responses and determine whether ignoring is an intermediate state (between aggression and retrieval) or a distinct behavioral state. Such a study will require a much larger sample size than was possible here, to opportunistically sample the natural variation in individual behavior.

## Conclusion

Aromatase expression in the male limbic brain is regulated with parental experience in an area-, hemisphere-, and experience-specific manner. Acquiring paternal pup-caring experience enhanced aromatase expression (number of aromatase positive cells) in brain areas processing olfactory information (PIR, CAM, MAM) and regulating aggression, sexual behavior and other social relations (MAM, VMH, LS). In centers essential for controlling parental behavior (MPOA, BNST, NAC), aromatase expression was instead present with left-hemisphere dominance and largely independent of experience with pups. In general, the experience-dependent changes in aromatase expression we observed in most brain areas did not mirror the previously reported changes in estrogen receptors (ERα) when pup-naïve males became pup-caring fathers.

## Acknowledgements

We thank H. Sabine Schmidt for assistance with animal care, immunocytochemistry, and microscopy. This study was supported by the Deutsche Forschungsgemeinschaft, EH53/19 to GE. PDG was funded by a postdoctoral fellowship from the Fonds Quebecois Recherche Nature et Technologie. PDG is currently supported by the Canada Research Chair Program. DAS was partially funded by a postdoctoral fellowship from the Kavli Neuroscience Discovery Institute.

## Abbreviations

BNST: bed nucleus of the stria terminalis
CAM: cortical amygdala
LS: lateral septum
LSD: lateral septum dorsal
LDI: lateral septum intermediate
LSV: lateral septum ventral
MAM: medial amygdala
MPOA: medial preoptic area
NAC: nucleus accumbens
PIR: piriform cortex
VMH: ventromedial nucleus of the hypothalamus.

