## Supplemental figures for "Brain aromatase and its relationship with parental experience and behavior in male mice"

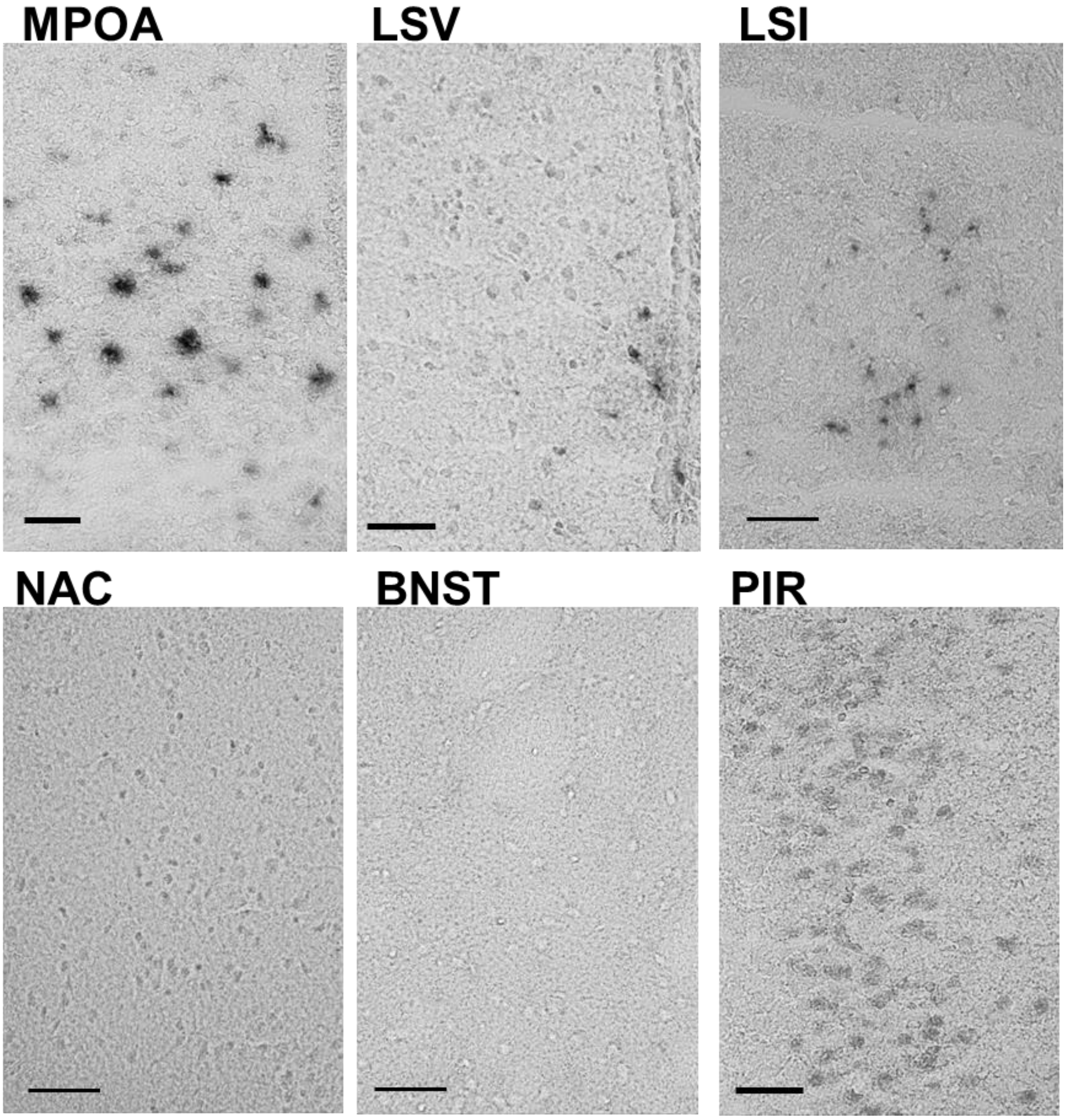


**Supplemental Figure 1.** Representative images of aromatase immunostaining in naïve males in the MPOA, LSV, LSI, NAC, BNST, and posterior PIR. Scale bar = 25 µm.

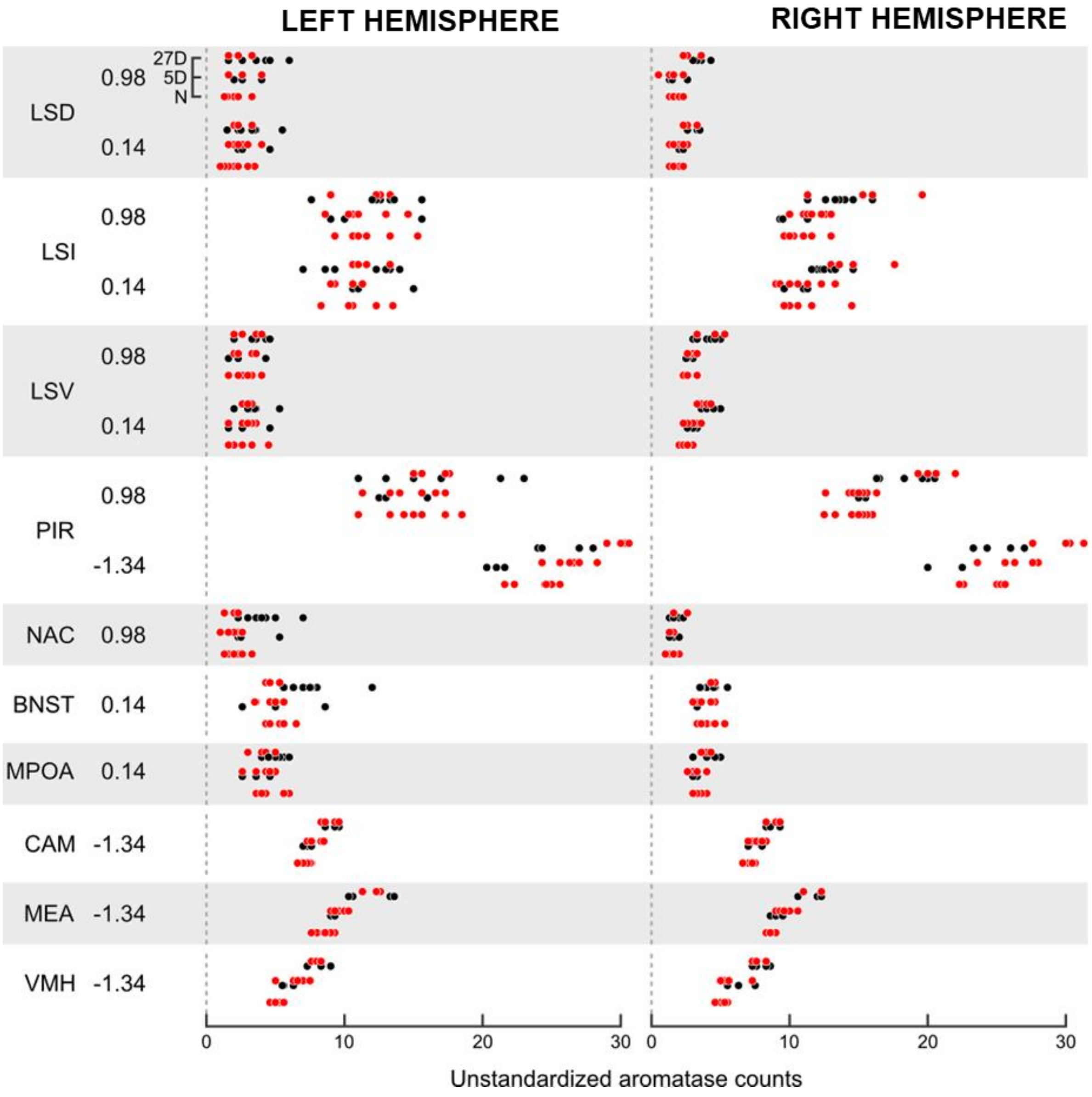


**Supplemental Figure 2.** Aromatase positive cell counts in the left and right hemisphere in various brain regions (including Bregma coordinates) in naïve males (N) and fathers with five days (5D) and 27 days (27D) of pup caring experience. Individual circles indicate individual males. The different colours (red or black) indicate the two cohorts of animals (see Methods for details).
